# Conservation of imprinted expression across genotypes is correlated with consistency of imprinting across endosperm development in maize

**DOI:** 10.1101/2024.08.16.608346

**Authors:** Kaitlin Higgins, Vital Nyabashi, Sarah Anderson

## Abstract

Imprinted expression is an essential process for seed viability affecting hundreds of genes in Zea mays endosperm, however most studies have examined just one time point for analysis. The focus on single time points can limit our ability to identify imprinted genes, and our ability to draw conclusions for the role of imprinting in endosperm. In this study we examine imprinted expression across four time points ranging from the transition to endoreduplication from mitotic division through the beginning of programmed cell death. Additionally, we assessed imprinting variation across eight diverse maize lines, six of which have never before been assessed for imprinting. Through this analysis we identify over 700 imprinted genes with varying consistency across time points including 258 genes imprinted at every time point and 104 genes displaying transient imprinting. We find a correlation between high consistency of imprinting across time and high conservation of parental bias across eight diverse maize lines reciprocally crossed with B73. Additionally, we identify evidence of imprinting for three zein genes that are critical for nutrient accumulation in the endosperm, suggesting that imprinting may play a more important role in seed composition than previously thought. Taken together, this study provides a more holistic view of imprinting variation across time and across genotypes in maize and enables us to more thoroughly investigate the complex imprinting landscape.

**Summary:** Though genomic imprinting is essential for seed development, changes in imprinted expression through endosperm development remain unclear. Here, the authors present a time series analysis of genomic imprinting in maize endosperm identifying over 1000 imprinted genes displaying consistent and transient imprinting. Additionally, the authors utilize imprinting data from B73 reciprocally crossed with eight diverse genotypes, and identify a correlation between consistency and conservation of imprinted expression. Together these results offer a more holistic view of imprinted expression in maize endosperm.

## Introduction

Endosperm is a tissue of angiosperm seeds responsible for providing nutrients for developing plants in the form of starch as well as oils and proteins. Additionally, endosperm is a rich source of carbohydrates that are consumed by humans and livestock. Endosperm is the result of double fertilization in flowering plants, and contains two copies of the maternal genome to one copy of the paternal genome. Before fertilization, the central cell undergoes demethylation of the maternal alleles through the activity of DNA-glycosylases active in the central cell and throughout endosperm development (Choi *et al*. 2002; Gutiérrez-Marcos *et al*. 2006; Gent *et al*. 2022). Together, triploidy and asymmetric demethylation create a unique transcriptomic environment. Due to this unique transcriptional environment, endosperm is the primary site of parent-of-origin dependent gene expression, which is known as genomic imprinting. Genomic imprinting is an essential process for endosperm development and disruption of imprinted expression results in inviable seeds (Scott *et al*. 1998; Stoute *et al*. 2012; Rebernig *et al*. 2015; Roth *et al*. 2019).

Though discovered over 70 years ago (Kermicle 1970), genomic imprinting has only been able to be studied at a genome wide level in the last 15 years. Prior to 2010, only a handful of imprinted genes had been identified due to rare phenotypic changes caused by mutations of imprinting regulators or genes themselves (Batista and Köhler 2020). More recently we have been able to identify hundreds of imprinted genes throughout the genome due to transcriptomic changes identified through RNA sequencing. From these studies we have determined imprinting occurs primarily in the endosperm of many flowering plants including Arabidopsis, maize, sorghum, castor bean, and rice (Haun and Springer 2008; Tiwari *et al*. 2010; Luo *et al*. 2011; Xu *et al*. 2014; Zhang *et al*. 2016; Yuan *et al*. 2017). While there is evidence of imprinted expression in various plants, just 10 genes have been identified as conserving imprinted expression between Arabidopsis and maize (Waters *et al*. 2013). Further efforts have been made to assess imprinted expression across individual lines within species and found that even within species imprinting is variable (Waters *et al*. 2013; Pignatta *et al*. 2014; Anderson *et al*. 2021).

Despite the dearth of conserved imprinted genes, these studies have also found features of imprinted genes that may be shared across flowering plants. Imprinted genes often overlap regions that are differentially methylated between endosperm and embryo (Gent *et al*. 2022) or are sites of asymmetric H3K27me3 silencing (Zhang *et al*. 2014; Moreno-Romero *et al*. 2016). These binary classes that often apply to imprinted genes have been further identified as predictors of the regulation of imprinting (Zhang *et al*. 2014). Additionally, the developmental pattern of expression for imprinted genes is often either endosperm specific or expressed in many tissues (Zhang *et al*. 2016).

Few studies have investigated imprinting across several different lines of the same species. One study investigated imprinted expression across five maize lines at 14 days after pollination (DAP), and concluded that only 11 MEGs and 24 PEGs were imprinted in all reciprocal crosses (Waters *et al*. 2013). Another focused on three Arabidopsis lines at 6 DAP and found 28 MEGs and 6 PEGs imprinted in all crosses (Pignatta *et al*. 2014). Even fewer studies have investigated imprinted expression across time, and looked primarily at embryonic imprinting (Meng *et al*. 2018), or focused on earlier endosperm development (Xin *et al*. 2013). Endosperm development is a dynamic process influenced by epigenetic regulation. At the beginning of endosperm development, the cells undergo replication through complete mitosis and cell division, and in maize this process lasts until 10 DAP. After 10 DAP, the endosperm shifts towards endoreduplication, or replication of chromosomes without cell division (Dai *et al*. 2021). At this point, the aleurone layer has differentiated from the main endospermic tissue, and starch accumulation has begun (Fath *et al*. 2000). At 16 DAP programmed cell death begins, concurrent with continuing protein and starch accumulation (Sabelli and Larkins 2009b), and by 28 days after pollination approximately half of the endosperm has undergone programmed cell death (Sabelli and Larkins 2009b). These shifts in endosperm development are accompanied by significant changes in expression, however many imprinting studies focus on one time point for analysis so we do not know how these changes impact imprinted expression.

In our present study we evaluate the transcriptome of endosperm from reciprocal crosses between B73 and W22 throughout development at four time points. These timepoints range from the beginning of starch accumulation and endoreduplication to the middle of the grain filling stage, allowing us to identify the transcriptional changes occurring during this significant transition. This experimental design also allows us to evaluate patterns of expression across time and identify differences in overall expression that could lead to imprinting, specifically whether an allele is permanently silenced or if it is transiently silenced, in order to tie observations to different regulatory implications. Additionally, we sequenced the transcriptomes of eight lines with full genome assemblies reciprocally crossed with B73 to see if there is a correlation between consistently imprinted genes and genes that display conserved imprinting across lines. In total, this study provides much more information about imprinted expression across development and genotypes of maize than has previously been available.

## Results

To investigate imprinted expression across time we performed reciprocal crosses between the maize lines B73 and W22 and harvested the ears at 11 days after pollination (DAP), 14 DAP, 17 DAP, and 21 DAP. These time points correspond to the transition to starch accumulation, prior imprinting studies, the beginning of programmed cell death, and progression of programmed cell death, respectively (Sabelli and Larkins 2009b). For each library, we dissected endosperm from 10 seeds off a single ear. For each time point, three biological replicates from each direction of the cross were collected, with the exception of 21 DAP, which had just 2 samples in the W22xB73 direction of the cross, for a total of 23 libraries. We then extracted RNA, reverse transcribed into cDNA, and performed Illumina sequencing. On average, we sequenced 45 million reads per library. To process the sequencing data, the reads were mapped to concatenated B73 and W22 genome assemblies, unique reads were counted, and imprinting status was determined with the reciprocal expression ration (RER) method (Anderson *et al*. 2021) which was designed to handle sequencing data from multiple assembled genomes. In parallel, we also mapped reads to a single consistent genome, B73, to assess global changes in endosperm expression across development without restricting to genes with sequence variants between the parental genomes. Correlation between pairs of samples was calculated and plotted, revealing correlation between samples at the same time point and in the same direction of the cross (Figure 1A).

**Figure 1:**
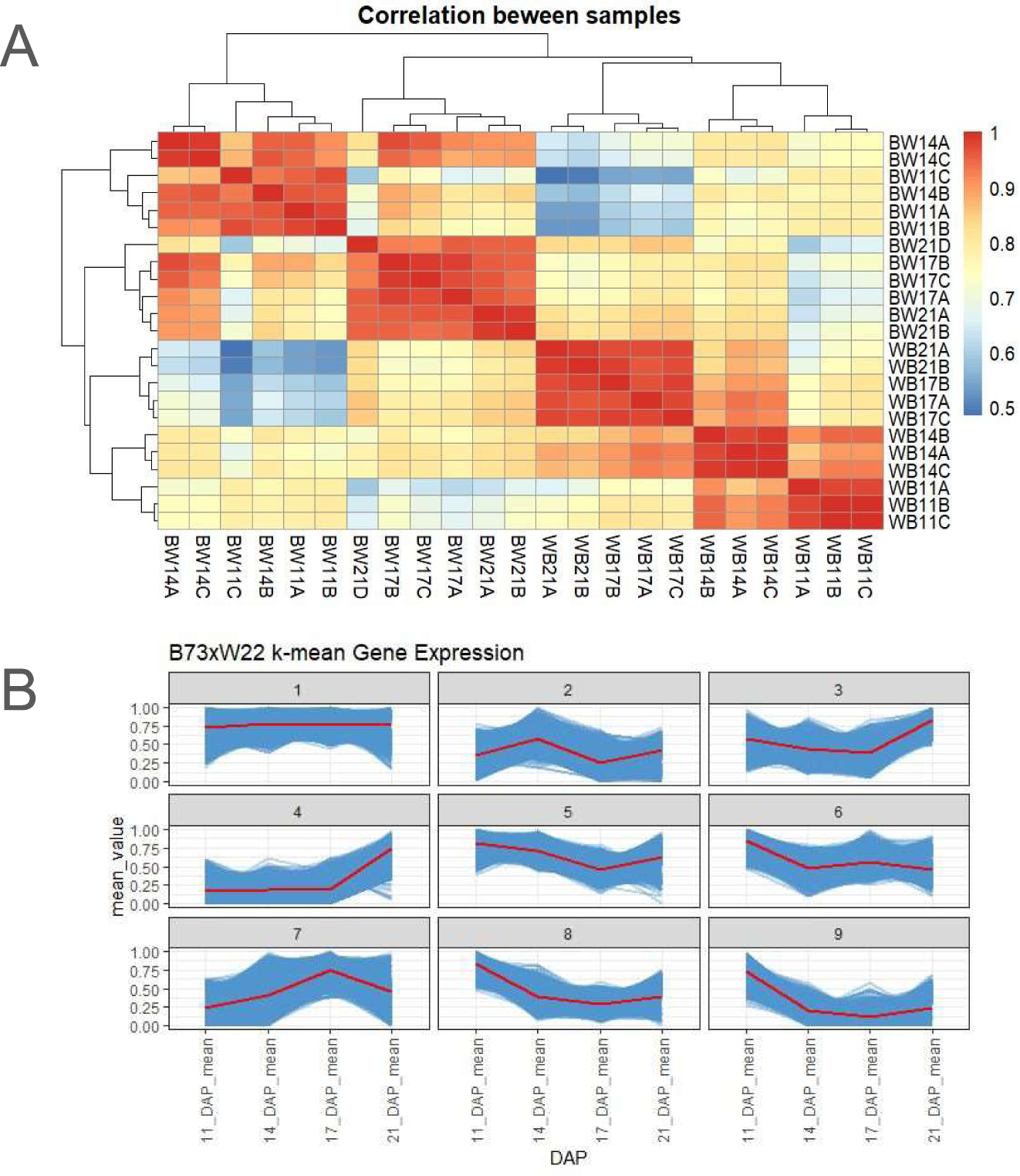
All gene expression correlation and peaks. (A) correlation plot ranging from 0.5 to 1 for all samples. (B) kmeans clusters of all genes in B73xW22 replicates over time. Reads for B73xW22 crosses were mapped to B73, then genes were normalized to rpm at each time point divided by max rpm at any time point. Blue lines represent average normalized expression across 3 replicates for each time point. Red lines represent the average of all genes in the cluster.

### K Means analysis of all time points mapped to B73

Expression changes across development can provide important insights into biological functions of tissues, thus our first step in this analysis was to identify major patterns of expression throughout development utilizing k-means clustering. This approach groups genes with similar expression together enabling us to see overall patterns. We chose for this analysis to be mapped to only one parental genome, B73, so that we could identify major patterns for all genes and identify features of genes in those clusters. Expression values were normalized by dividing the expression at each timepoint by the maximum expression at any time point, resulting in expression for each gene normalized to a scale from 0 to 1. Using the samples where B73 was the maternal parent (BxW) we identified nine clusters of expression across all four timepoints. Among these, we saw clusters where expression peaked at each timepoint (Figure 1B), clusters that consistently increase, and clusters with genes that are generally consistent across time. To validate the consistency of expression clusters across the reciprocal direction of the cross, we mapped W22 maternal (WxB) samples to the B73 genome and placed them in the same clusters identified in BxW and found that the expression changes were largely consistent in both directions of the crosses (Figure S1A). The maize prolamin gene family, also known as zeins, is known to account for nearly 65% of all transcripts found in endosperm (Li and Song 2020), making their expression an easily identifiable benchmark for expression pattern analysis. We found that zeins inherited maternally (BxW) or paternally (WxB) followed similar patterns in the clusters to which they were assigned (Figure S1B). We therefore chose to focus the following DE analysis on the time-point analysis of all genes in only BxW (B73 maternal) crosses.

### DE expression in relation to earliest time-point

To define a core set of dynamically differentially expressed genes across our time series, we next evaluated differential expression across time in comparison to the earliest time point of 11 DAP. Using the R package DESeq2 (Love *et al*. 2014), we called differential expression (log2FC >1, FDR < 0.05) for genes between 11 DAP and each later time point. We found 2000-3300 differentially expressed genes (DEGs) both up and down regulated at each of the three later timepoints in comparison to 11 DAP (Figure 2A). Next, we compared the sets of genes to find which are continuously DE and which are timepoint specific (Figure 2B). We found that 30% of upregulated genes and 24% of downregulated genes are consistently DE across all three time points when compared to 11 DAP. Of the genes that are not DE at all timepoints, we find 43% (1,846) of upregulated genes are DE at just one of the three compared times, while 47% (2,130) of downregulated genes are DE at just one time. Nearly half (23%, 1,051) of the genes downregulated at one time point, are down regulated at 17 DAP (Figure 2C), which coincides with the time at which zeins are being expressed at the highest rates (Figure S1B).

**Figure 2:**
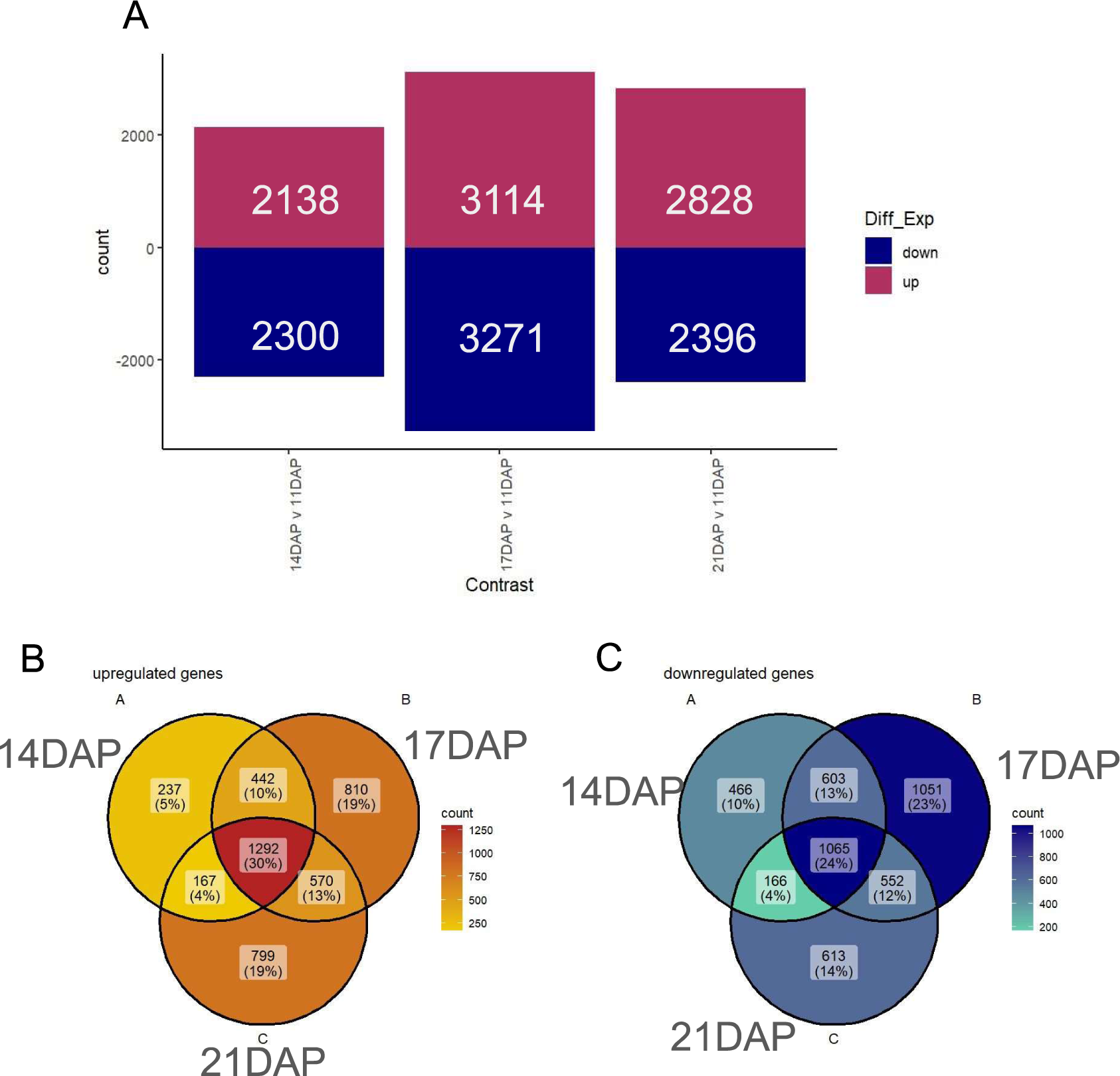
Differentially expressed genes over time. (A) DE genes at each time point in relation to 11DAP, upregulated genes represented in maroon and downregulated genes represented in navy. (B) Venn Diagram ofupregulated DE genes at each timepoint to highlight genes that are DE at more than one time point. (C) Venn Diagram of downregulated DE genes at each timepoint to highlight genes that are DE at more than one time point.

### Imprinting over time

One expression pattern we sought to investigate further is genomic imprinting as this is an essential pathway for endosperm development (Batista and Köhler 2020), yet the dynamics of imprinting across endosperm development are still not fully understood. To evaluate imprinting across time, we first mapped all reads to concatenated genomes removing multi-mapping reads. We then called imprinted expression for each gene by applying DESeq2 to reads in contrasting directions of reciprocal crosses (log2FC >= 1, FDR < 0.05). Genes were defined as imprinted only when significantly more than 2-fold differentially expressed when inherited maternally versus paternally in order to account for differential contributions of maternal and paternal genomes to chromosome numbers in the triploid endosperm as in (Anderson et al 2021). We analyzed imprinted expression at each timepoint and found over 200 MEGs and over 50 PEGs at each timepoint and in each inbred line (Figure 3A). We then compared imprinting calls at each timepoint to determine which genes display variable imprinting across time, and which are imprinted at all time points. We found 282 (39%) of all MEGs and 108 (33%) of all PEGs identified in this experiment are consistently imprinted (Figure 3B). Our stringent statistical cutoffs for imprinting can result in under-calling imprinted expression, so we further assessed genes for parentally biased expression.

**Figure 3:**
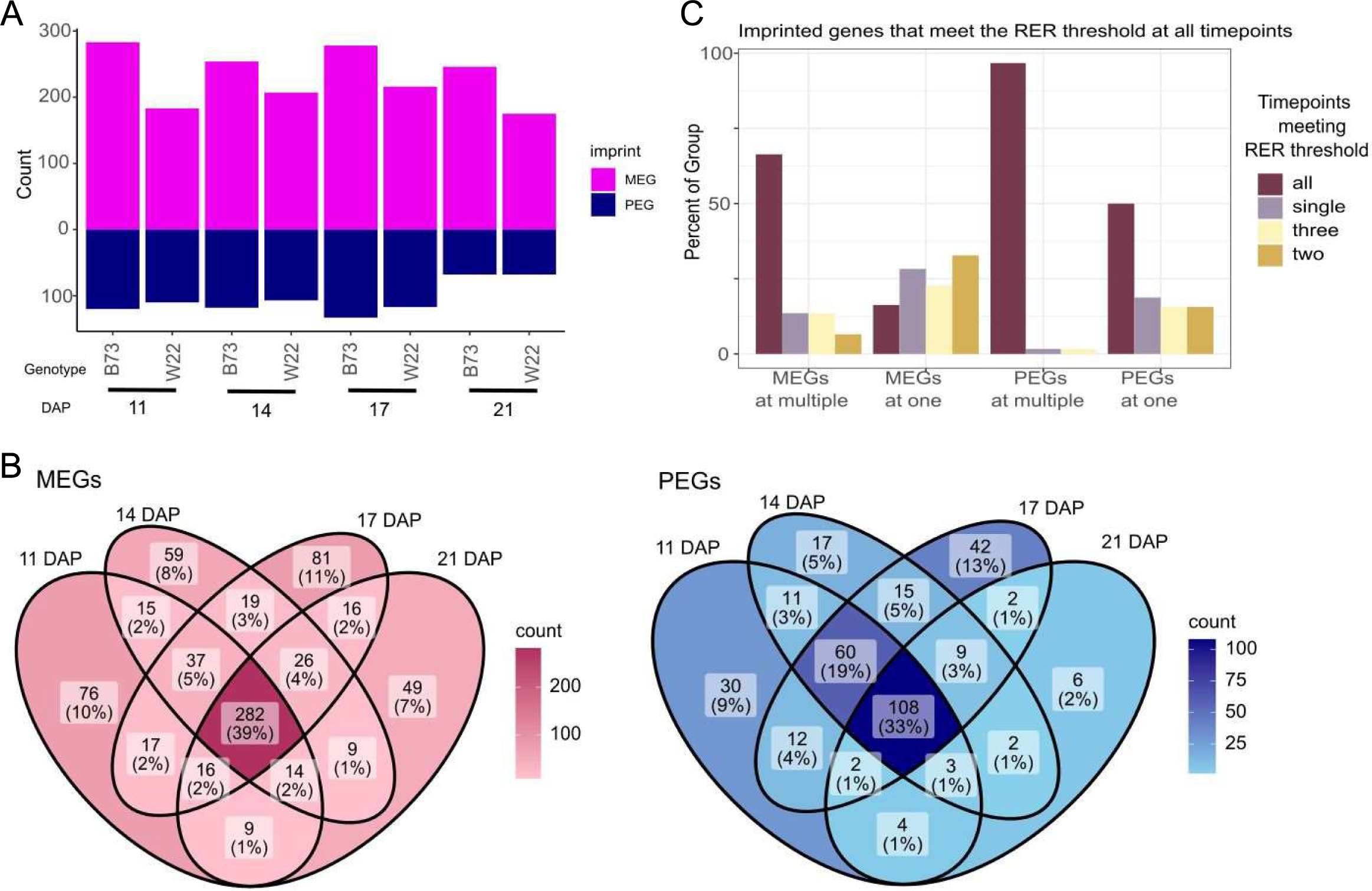
Identification of imprinted genes. (A) Imprinted gene count. Imprinted genes identified at each timepoint in each genotype of the cross, purple indicates maternally expressed genes (MEGs) navy indicates PEGs. (B)Venn diagrams ofMEGs and PEGs comparing genes imprinted at each timepoint. Dark blue and red indicate the highest number of imprinted genes occur in that segment of the venn diagram. (C) Frequency of genes meeting RER threshold at multiple or single time points when not meeting imprinting cutoffs at all timepoints. Colored by the number of times they reach the RER threshold.

To define parentally biased expression, we utilized cutoffs based on reciprocal expression ratio (RER). RER is a measure of the normalized expression when inherited maternally versus paternally. It is calculated by dividing the reads per million (RPM) when inherited maternally by the sum of maternal and paternal RPM. This results in a value between 0 and 1, where 1 indicates all transcripts are from the maternal allele and 0 indicates all transcripts are from the paternal allele. Thus, RER correlates to maternal preference, or the percentage of reads expressed from the maternal allele (Anderson *et al*. 2021). Biparentally expressed genes have an RER around 0.66 due to the presence of two copies of the maternal genome to the one copy of the paternal genome. Significant deviance from 0.66 would indicate parentally biased expression with high RER indicating maternal preference and low RER indicating paternal preference. In accordance with Anderson et al 2021, we identified imprinted expression using RER cutoffs of 0.9 for MEGs, and 0.3 for PEGs, however for biased expression, we used less stringent cutoffs of 0.4 for paternal bias, and 0.8 for maternal bias.

We grouped the imprinted genes based on whether they were imprinted at one time point or multiple timepoints. Next, we evaluated whether these groups often met the RER thresholds to see if there were transient imprints that generally display parentally biased expression but do not meet the stringent thresholds to be defined as imprinted. To do this, we calculated the RER for each timepoint for each gene, then counted how many timepoints at which the RER met the minimum threshold to be defined as parentally biased (single, two, three, or all). We found that 66% of genes defined as MEGs at multiple timepoints had RERs meeting parentally biased expression at all timepoints, whereas just 16% of MEGs at a single time point had RERs meeting that threshold. Additionally, 99% of PEGs at multiple timepoints had RERs meeting the parental bias threshold, and 50% of PEGs at single time points did as well (Figure 3C). This indicates that there may be allelic bias present even when the expression data do not meet the strict cut offs used to call imprinted expression, so further classification was performed to distinguish true dynamic imprinting rather than apparent dynamics caused by stringent cutoffs.

### Imprinting patterns analysis

The high percentage of MEGs and PEGs that met parentally biased thresholds when not imprinted suggests more nuance to imprinted expression than strict cutoffs for calling imprinting can discern. To evaluate this nuance while accounting for allelic bias that may not rise to the threshold of imprinted expression, we assessed allelic patterns for imprinted genes over time. We removed all genes called as imprinted that are never expressed above 1 rpm to focus on high confidence imprinted expression. A total of 338 genes identified as imprinted did not meet this threshold leaving us with 710 high confidence imprinted genes. We then categorized these into four broad categories. The first category (Group 1, 258 genes) contains genes expressed above 1 rpm at all timepoints and imprinted in the same direction at all timepoints. The second category (Group 2, 42 genes) contains genes that are not expressed at all timepoints, but are imprinted in the same direction at all time points at which they are expressed above 1 rpm. The third category (Group 3, 306 genes) contains genes that are imprinted at one time point or more, but also display parental bias in the same direction as the imprint when not imprinted. Finally the last category (Group 4, 104 genes) contains the set of genes with truly variable expression. In order to make this call, a stricter expression cutoff of 5 rpm in addition to variable imprinting calls was implemented. After creating all four groups, 123 genes meeting the 1 rpm expression threshold were not assigned to any group (Figure 4A).

**Figure 4:**
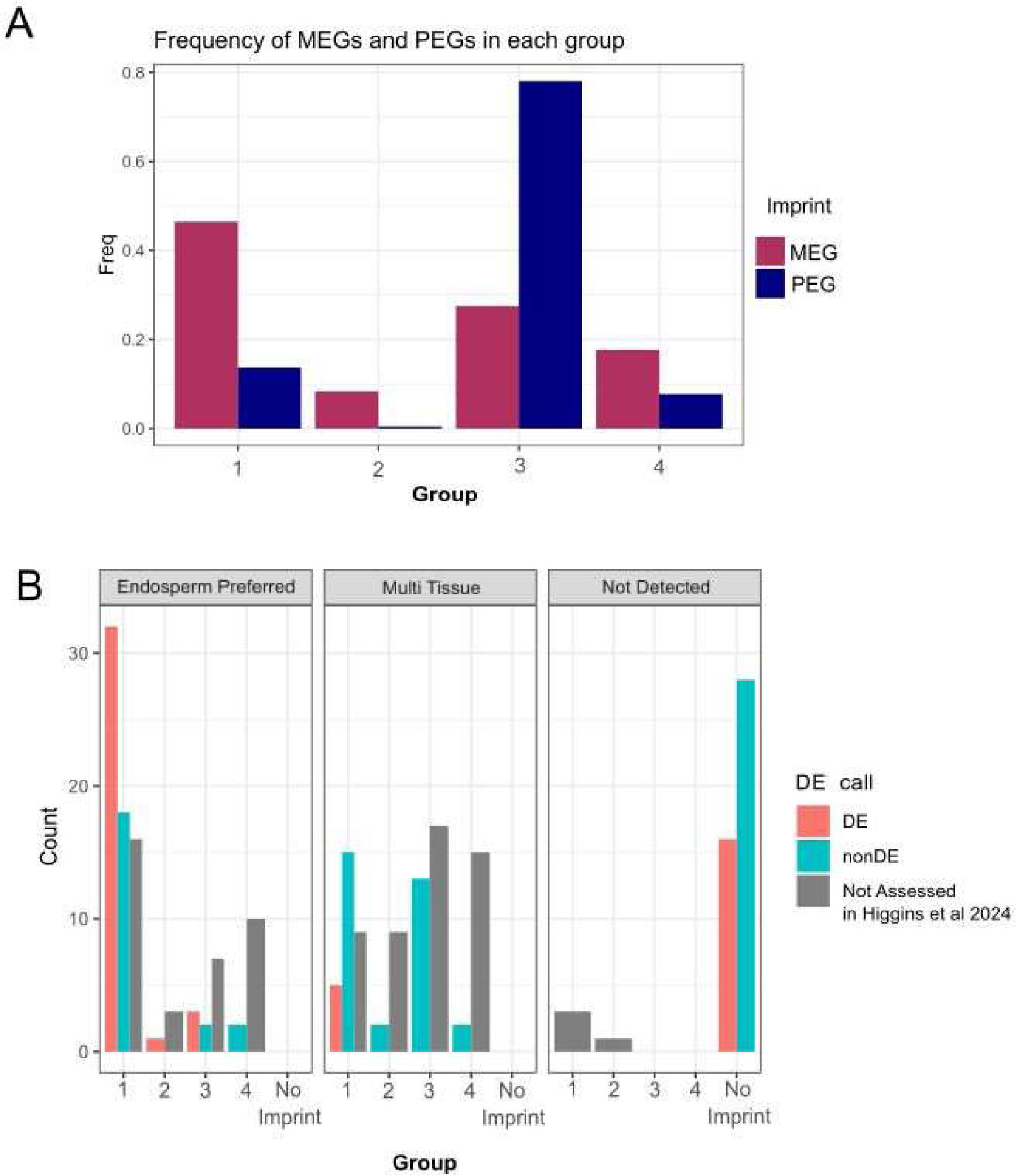
Grouped imprinted genes. (A) Frequency ofMEGs and PEGs in each of the four expression groups. MEGs in burgundy, and PEGs in navy blue. (B) MEGs with expression in Mdrl study detailed in chapter 2 by group and tissue specific expression. Red indicates differential expression in the mutant, and thus regulation by Mdrl. Blue indicates non-differential expression in the mutant, which could be either redundant regulation by Mdrl or no Mdrl involvement in regulation. Grey indicates the gene was not assessed in Higgins *et. al* 2024. NA category of tissue specific expression indicates it could not be assessed in the W22 expression atlas.

We found that 228 MEGs (46.4%), and 30 PEGs (13.6%) belonged to group 1. This shows that nearly half of MEGs meet the thresholds used to determine imprinting consistently across time, while in contrast just 13% of PEGs are consistently imprinted. In group 2 we find an additional 41 MEGs (8%), and 1 PEG (0.4%) with consistent imprinting whenever expressed. Group 3 contains 151 PEGs (75.8%), and 101 MEGs (24.8%) with consistently biased expression that lack statistical support in at least one time point. Finally, group 4 contains 17 PEGs (8.5%) and 87 MEGs (21.3%) with truly variable imprinting. These groups can be broadly described in terms of consistency where Group 1 would be the most consistently imprinted genes, and genes in Group 4 would display inconsistent or transient imprinting.

Imprinted expression originates from multiple different mechanisms. Broadly, four groups of imprinting have been proposed by categorizing MEGs and PEGs based on whether expression is endosperm preferred or expressed in multiple tissues across plant development (Zhang *et al*. 2014; Batista and Köhler 2020). The best understood mechanism, maternally activated expression, occurs when the maternal alleles are demethylated by DNA glycosylases in the central cell before fertilization, allowing for imprinting and expression only in endosperm tissues since the paternal allele is never expressed. One example of this regulatory pathway is *Fie1* which is imprinted and regulated by the demethylase activity of Demeter in Arabidopsis (Hsieh *et al*. 2009). To investigate if there is an association between imprinting across time and expression across development (Monnahan *et al*. 2020), we evaluated our groups for endosperm preferred expression versus multi-tissue expression. As expected for maternally activated MEGs, the majority of Group 1 MEGs that are expressed predominantly in the endosperm are regulated by DNA glycosylase *Mdr1* (Figure 4B, (Higgins *et al*. 2024)).

### Conservation of imprinting across genotypes of maize

Imprinting has previously been shown to be dynamic with some alleles showing imprinted expression while others are not imprintable (Waters *et al*. 2013). Due to the dynamic nature of imprinting, we sought to investigate the relationship between variable imprinting across genotype and variable imprinting across time. In an effort to identify the variability of imprinting within *Zea mays* at greater depth, we utilized previously published sequencing data as well as generated new sequencing data from reciprocal crosses between B73 and other lines with published genome assemblies. Previously published data included sequencing data for B73 reciprocally crossed with Ki11 and Oh43 from Waters et al 2013, and W22 reciprocally crossed with B73 from Anderson et al 2021 and analyzed at 14 DAP. Newly acquired sequencing data includes the lines NC358, CML333, M162W, B97, Ky21, and Oh43 reciprocally crossed with B73 and dissected then analyzed at 14 DAP. On average, we sequenced 14 million reads per library. For all libraries, RNA-seq reads were mapped to concatenated genomes of B73v5 and the respective alternate parent, unique reads were counted, and RER was calculated to quantify parent-of-origin biased expression (Anderson *et al*. 2021). The total number of imprinted genes is highly variable across lines (Figure 5A), while the RER counts are more consistent (Figure 5B) and may help us identify the genes that are required for seed viability when imprinted or biased in the endosperm across multiple lines, like *Fie1. Fie1* is an imprinted gene required for endosperm development (Hermon *et al*. 2007).The total count for all parentally biased genes using RER ranged from 2500-5000 for maternal bias (minimum 0.8 RER), and from 2000 to 3000 for paternal bias (maximum 0.3 RER) (Figure 5B). Syntologous, single-copy genes across maize genomes have been previously called (Woodhouse *et al*. 2021), which we utilized to identify syntelogs in the surveyed genomes. We then plotted the parental bias across lines for genes within each time series group to identify potential relationships (Figure S2A). *Fie1*, as we might expect, falls into Group 1 MEGs, or MEGs consistently imprinted across time, and when we evaluate RER across lines we see that out of 8 reciprocal crosses, the *Fie1* RER meets the cutoff the majority of the time (Figure 5C, Figure S2B).

**Figure 5:**
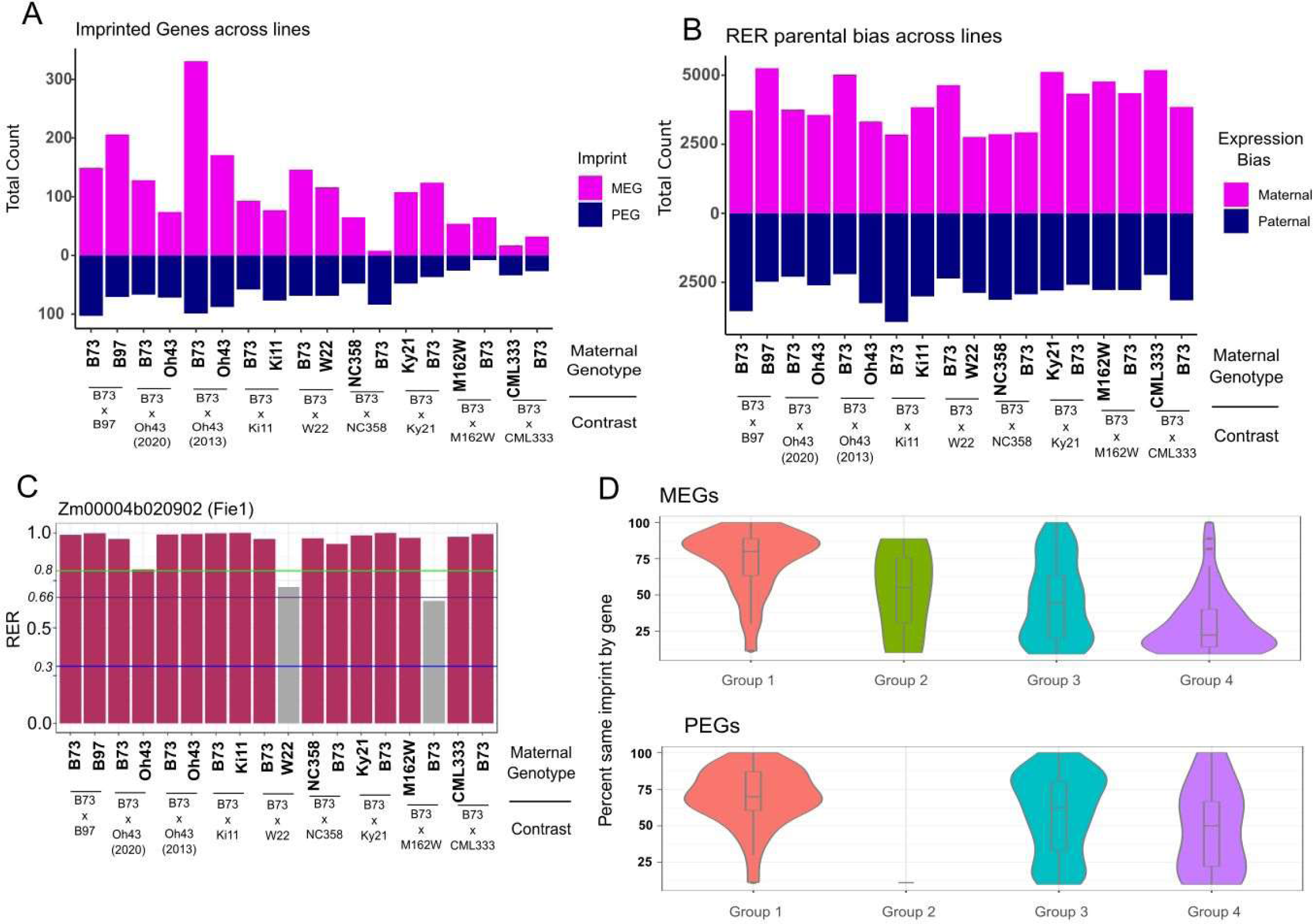
Imprinted expression at 14 DAP across maize genotypes. (A) Genes meeting imprinting thresholds for maternally expressed genes (pink) and paternally expressed genes (navy blue) in each direction of each reciprocal cross. (B) Genes meeting reciprocal expression ratio for paternally biased expression (navy blue) or maternally biased expression (pink) for each direction of each reciprocal cross. (C) Barplot of Fiel maternal preference across lines. Maroon indicates the maternal preference in that cross meets the threshold for RER to be maternally expressed, Grey indicates biparental expression. Purple horizontal line indicates biparental expression average (0.66), green horizontal line indicates maternal RER threshold (0.8), blue horizontal line indicates paternal RER threshold (0.3). (D) Rates of imprinted MEGs and PEGs in the four expression groups maintaining parental bias in reciprocals across genotypes. These are limited to genes present in at least four genotypes.

To evaluate any correlation between the previously defined groups of imprinted genes and consistency across lines, we compared the imprinted genes in Groups 1, 2, 3, and 4 to parentally biased expression across lines. We first calculated the percent same imprint by gene, which is the total number of genotypes in which the gene is parentally biased divided by the number of genotypes with an expressed syntelog. This measure is limited to only genes with informative genetic variants present between alleles in reciprocal crosses, since genes with identical sequence could not be assessed and were not included in the comparison. Next, we plotted genes present in at least 4 genotypes by group and imprint (MEG or PEG) against the percent same imprint by gene measure to determine if there were any trends by group (Figure 5D). We found that the mean percentage of conserved imprinting across lines goes down in a stepwise manner across MEGs in each group, with Group 1 MEGs maintaining imprinting across ∼75% of lines in which they were identified, down to ∼25% of Group 4 MEGs maintaining imprinting. The decrease in conservation across lines corresponds with a decrease in the consistency of imprinting across development, as Group 1 are the most consistently imprinted group, and Group 4 is the group that displays transient imprinting. This decrease in imprinting conservation across lines is also evident among PEGs, however group 4 PEGs still display conserved imprinting at a higher rate than Group 4 MEGs.

### Imprinted zeins

While investigating the total expression of imprinted genes, we found that a handful of MEGs had high expression at all time points. To evaluate them further we looked at protein annotations for W22 and B73 from maizeGDB that are associated with these genes and found that three (Zm00004b020603, Zm00004b020785, Zm00004b020789) are associated with alpha-zeins. Zeins are starch accumulation proteins that are highly expressed in late endosperm development, and are the main storage protein for endosperm (Li and Song 2020). Further, when we blasted the sequence of Zm00004b020603 against the B73v5 genome, we found homology to floury2 (Zm00001eb170070) along with homology between flanking genes of floury2 in both genomes, which we then evaluated using CoGe (Lyons and Freeling 2008); Figure 6). This synteny indicates that Zm00004b020603 is the ortholog of floury 2 in W22, and is imprinted in this genotype. The reason for this particular version of floury2 displaying imprinted expression is not clear as they both have upstream long terminal repeat transposons, and there is no clear sequence difference in the promoter regions. These results identify the W22 homolog of floury2, and provide further evidence that zeins can be imprinted in some genotypes.

**Figure 6:**
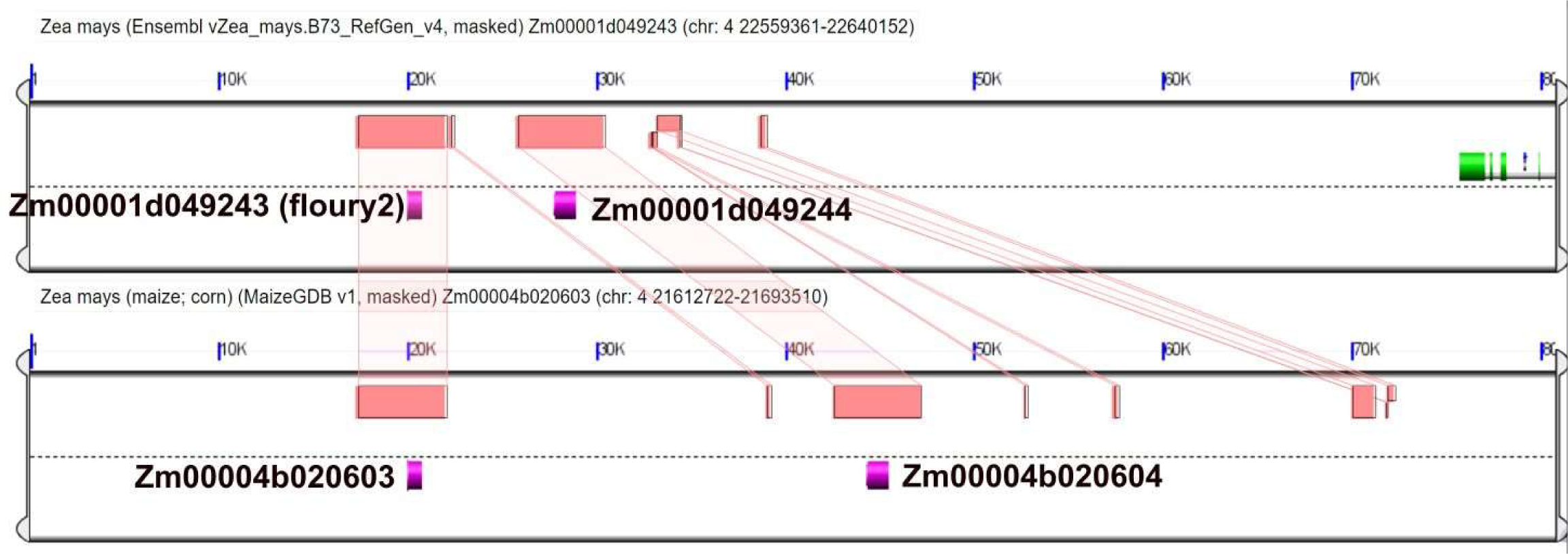
Floury2 W22 allele synteny. Comparison of syntenic regions (red boxes) between floury2 in B73 (Zm00001d049243) and Zm00004b020603 rendered using CoGe online tool. Including nearby genes that share synteny.

## Discussion

To better understand imprinting dynamics, in this study we utilized four time points reaching from the beginning of endoreduplication to mid grain filling stage (Sabelli and Larkins 2009a) in combination with the RER method of imprinting detection which allows for evaluation of imprinting in genes that display presence absence variation (Anderson *et al*. 2021). We found that each time point assessed had 190-290 MEGs and 50-110 PEGs in each direction of the cross totalling 1048 unique genes across all time points. Due to strict cutoffs for identifying imprinted expression, some of these genes may still display parentally biased expression when not imprinted. After categorizing imprinted genes into four groups based on the extent of imprinted and parentally biased expression over the time points, we found that around 40% of MEGs meeting our expression threshold of at least 1 rpm at every time point were imprinted at every time point, while just 15% of PEGs were imprinted at every time point. Previous studies have found that well known imprinted genes *Fie1* and *Mez1*, which are important for the FIS-PRC2 complex, appear imprinted consistently up to 15 DAP (Xin *et al*. 2013). Our study also found *Fie1* to be imprinted at all time points up to 21 DAP, however *Mez1* fell into Group 3, which displays parentally biased expression at every time point but does not meet the threshold requirements to be called imprinted. The shift to maternally biased expression occurred later in development than Xin et al surveyed in 2013, and thus these results are still consistent with what the authors found. Additionally, we identified 104 imprinted genes (87 MEGs, 17 PEGs) that display truly variable imprinting in that they are expressed above 5 rpm at every time point, and lose or gain imprinted or parentally biased expression. These results highlight the importance of studying multiple time points due to changes in gene expression through endosperm development.

After classifying imprinted genes into expression groups, we were then able to utilize tissue expression data to associate the consistency of imprinted expression with the potential method of regulation. Tissue expression is one factor useful for predicting the type of imprinting regulation, for example, endosperm specific expression in MEGs is associated with maternal activation of imprinting through demethylation of the maternal alleles (Zhang *et al*. 2014; Batista and Köhler 2020). We saw that Group 1 MEGs contained the highest proportion of genes with endosperm preferred expression, which suggests that maternal activation predominantly leads to consistent expression of maternal alleles through endosperm development.

Utilizing a panel of reciprocal crosses with B73 and other maize genotypes with full genome assemblies, we were able to establish a correlation between consistent imprinting through endosperm development and conservation of parentally biased expression across genetic lines. Through this analysis we find that genes displaying consistent imprinted expression from 11-21 DAP are more correlated with conserved imprinted expression at 14 DAP across genetic lines in which they are present. We also find that genes displaying transient imprinting are less correlated with conserved imprinted expression across lines at 14 DAP. Maternally expressed genes show a more dramatic difference between consistent and transient imprint than paternally expressed genes, which is likely due to the higher rates of presence absence variation among MEGs. This information could be further utilized to identify causative factors involved in imprinting regulation in depth.

Finally, through this study we were able to identify maternally expressed imprinted zeins. Zeins generally provide a substantial portion of endosperm proteins, and are among the most highly expressed endosperm genes. This finding supports the kinship hypothesis (Feil and Berger 2007) which suggests that imprinting occurs as a way for parental genomes to influence nutrient allocation to offspring for higher fitness. Interestingly, these imprinted zeins originated from W22 and did not show a substantial difference in sequence or location of nearby transposable elements from their B73 syntelogs. This lack of substantial change between syntelogs could be utilized to understand imprinting regulation through further experimentation and analysis.

In conclusion, imprinted genes are most often either consistently imprinted or maintaining a parental bias over endosperm development at the time points assessed. Transient imprinting does occur, but is less common and not well understood. We additionally found a correlation between consistency of imprinting over time and conservation of parentally biased expression across maize genotypes as well as a difference in imprinted expression between two highly similar zein genes. Together these findings can be utilized to find sequence variants that influence imprinted expression and further understand imprinting regulation.

## Materials and Methods

### Reciprocal time series RNA-sequencing and tissue collection

The maize inbred lines W22 and B73 were grown in Ames, IA during the summer of 2023. These lines were reciprocally crossed then harvested at 11, 14, 17, and 21 days after pollination within 40 minutes of the time they were pollinated. The maize inbred lines B73, B97, and Oh43 were grown in Ames, IA during the summer of 2020, and maize lines B73, Ky21, NC358, CML333, and M162W were grown in Ames, IA during the summer of 2022. In each year B73 was reciprocally crossed with the other inbred lines and harvested 14 days after pollination Manual dissection of approximately 10 kernels from each ear were pooled into one biological replicate. Raw reads for Ki11 reciprocals were obtained from the NCBI bioproject archive PRJNA148441 (Waters *et al*. 12/2011).

RNA was extracted from each of three biological replicates of reciprocals at each time point using the Qiagen RNeasy Mini Kit (cat # 74104) after pulverizing wet endosperm in liquid nitrogen. Sample quality checking was performed using an Agilent Bioanalyzer 2100, prior to diluting each sample to 100ng/uL for library preparation and sequencing submission. Paired-end cDNA libraries were prepared using the NEBNext Ultra II Directional RNA kit (cat #E7760S). Samples were then sequenced on the Illumina NovaSeq 6000 using 150 paired-end sequencing at the Iowa State DNA facility, resulting in an average of 45 million reads per sample. Sequence reads were trimmed using cutadapt (Marcel 2011) (parameters -a AGATCGGAAGAGCACACGTCTGAACTCCAGTCAC -A AGATCGGAAGAGCGTCGTGTAGGGAAAGAGTGTAGATCTCGGTGGTCGCCGTATCATT -m 30 -q 10 --quality-base=33) then aligned to the concatenated parental genome assemblies. Concatenated parental genome assemblies were created by appending the W22-NRGene assembled chromosomes 1-10 (Springer *et al*. 2018) to the B73v5 assembled chromosomes 1-10 (Woodhouse *et al*. 2021) to create a single fasta file. The time series samples were additionally aligned in parallel with aligning to the individual B73v5 genome using hisat2 (Kim *et al*. 2019) (parameters -p 6 -k 20). Counts were determined through htseq-count (Anders *et al*. 2014) (parameters -f bam -r pos -s no -t all -i sequence_feature -m union -a 0) using a gene annotation file created by concatenating the B73v5 and W22 annotation files (Woodhouse *et al*. 2021). The counts were then normalized to reads per million (rpm). Differential expression was determined using R packages, and only features with |>1| Log2FoldChange along with an FDR padj < .05 were called as differentially expressed.

### Alignment to B73 of time series samples

Using the previously sequenced and trimmed samples, reads were aligned to the B73v5 genome assembly using hisat2 (Kim *et al*. 2019) (parameters -p 6 -k 20). Counts were determined through featureCounts (Liao *et al*. 2014) (parameters -p -O -t gene -g ID) using genes from B73v5 annotation files on MaizeGDB (Woodhouse *et al*. 2021). Counts were grouped based on maternal parent to create count tables. The counts were then normalized to reads per million(rpm). Differential expression was determined using the R package DESeq2 (Love *et al*. 2014), and only features with |>1| Log2FoldChange along with an FDR padj <.05 were called as differentially expressed.

### Imprinting calculation

To estimate parent of origin bias for expression and to call imprinting, the RER method was implemented (Anderson *et al*., 2021). Briefly, count tables including both directions of reciprocal crossed endosperm were analyzed by DESeq2 to identify genes with significantly more than 2-fold differences in expression when inherited maternally vs paternally using parameters lfcThreshold = 1 and altHypothesis = “greaterAbs”. RER was then calculated using RPM normalized count tables by dividing the gene rpm when inherited maternally by the sum of gene rpm when inherited maternally plus paternally.

### K means clustering

To evaluate the fluctuation in gene expression across timepoints, k means clustering was performed separately for each direction of the reciprocal crosses of B73 and W22 mapped to B73v5. First, rpm values were averaged by time point then divided by the maximum value for each gene. This created values for each gene at each timepoint ranging from 0-1 where 1 is the highest value for normalized expression. The R package stats (R Foundation for Statistical Computing 2018) was then used to cluster all expressed genes into one of nine clusters based on the change in expression over time. To test for consistency across reciprocal crosses, cluster assignments from the B73xW22 crosses were extracted and the expression patterns for W22xB73 libraries were plotted (Fig S1A).

### Feature Assessment for Imprinted Genes

#### Group assignments

Genes identified as imprinted at any time point were organized into groups in order to describe consistent vs transient imprinting through seed development. In order to ensure high confidence calls, genes were first filtered by expression, requiring a mean of 1 rpm across all time points. These were then subdivided into groups based on imprinting patterns. Group 1 required genes be imprinted and expressed at all time points assessed. Group 2 required genes be imprinted at any time point in which they were expressed, however they did not have to be expressed at all time points. Group 3 consisted of genes that were parentally biased when not imprinted. For the last portion of imprinted genes, a stricter rpm cutoff of 5 rpm was applied so that Group 4 consists of clearly expressed genes that are imprinted at at least one time point but either gain or lose imprinted expression.

#### Tissue expression pattern assignments

Utilizing a W22 expression atlas (Monnahan *et al*. 2020), we evaluated the expressed W22 genes for endosperm preferred expression or multi-tissue expression. Briefly, RPM was calculated for all genes across 10 tissues and defined as endosperm preferred when more than 65% of all reads originated in the endosperm replicates and multi-tissue when expressed more broadly. Genes were then grouped according to imprinting group and direction of imprinting or biparental expression. A list of DE genes between WT W22 and *mdr1* mutant W22 (Higgins *et al*. 2024) was then compared to the tissue specific expression and expression groups assigned to W22 imprinted genes identified in this study.

#### Identification of syntologous genes across lines

To identify syntologous genes across lines, the archived MaizeGDB_Maize_pangene_2020_08.tsv file (https://download.maizegdb.org/Pan-genes/archive/pan-zea-2020/MaizeGDB_Maize_pangene_2020_08.tsv) with the script format_geneIDs.pl found at https://github.com/kmhiggins/Imprinting_timeseries_2024 to select syntelogs for just the genomes assessed in this study. The resulting file was then read into R where it was filtered for single gene syntelogs between B73 and opposite parent genotypes. Syntologous genes were matched to previously calculated RER and imprinting calls and ggplot2 (Wickham 2009) was utilized to visualize comparisons of syntelogs across genotypes.

#### Identification of floury2 in W22

Imprinted zeins were identified through protein annotations for W22 and B73v5 available at maizegdb.org (Woodhouse *et al*. 2021). W22 zein genes were then blasted to B73v5 for comparison, and one, Zm00004b020603, was found to be similar to floury2 in B73. The W22 and presumed syntelog B73 genes were then input into CoGe (Lyons and Freeling 2008) GEvo tool. B73v5 was unavailable through this tool so the B73v4 version of floury2 was used for comparison instead. For Zm00001d049243, genome parameters were B73_RefGen_v4.41.gff3. For Zm00004b020603, genome parameters were MaizeGDB_zea_maysw22_2.0_core_53_87_12_chr_scaffold.v2.gff. Parameters were changed to yes for the following results visualization options: color anchor gene yellow, draw all HSPs on top, Draw all HSPs on same track, Color features overlapped by HSPs, Show Other Features, Show pre-annotated CNSs, Show pre-annotated Gene spaces. All other parameters remained in default setting. Original analysis can be viewed at https://genomevolution.org/r/1s2m9

## Acknowledgements

We would like to thank the Iowa State DNA facility for their sequencing services and Conner Valentine for his contribution to RNA extraction.

## Data Availability

All raw sequencing data generated in this study have been submitted to the NCBI BioProject database (https://www.ncbi.nlm.nih.gov/bioproject/) under accession numbers PRJNA1123772 (8 genotype reciprocals) and PRJNA1110585 (time series).

All scripts used to define DE genes, imprinted genes, features of genes, and to generate figures can be found in the github repository at https://github.com/kmhiggins/Imprinting_timeseries_2024.

## Author Contributions

SNA and KMH designed the research, KMH and VIN generated the data. KMH and VIN analyzed data. KMH and SNA wrote the paper with input from all authors.

