## Supplemental Figures for "Conservation of imprinted expression across genotypes is correlated with consistency of imprinting across endosperm development in maize"

### Supplemental Figure 1

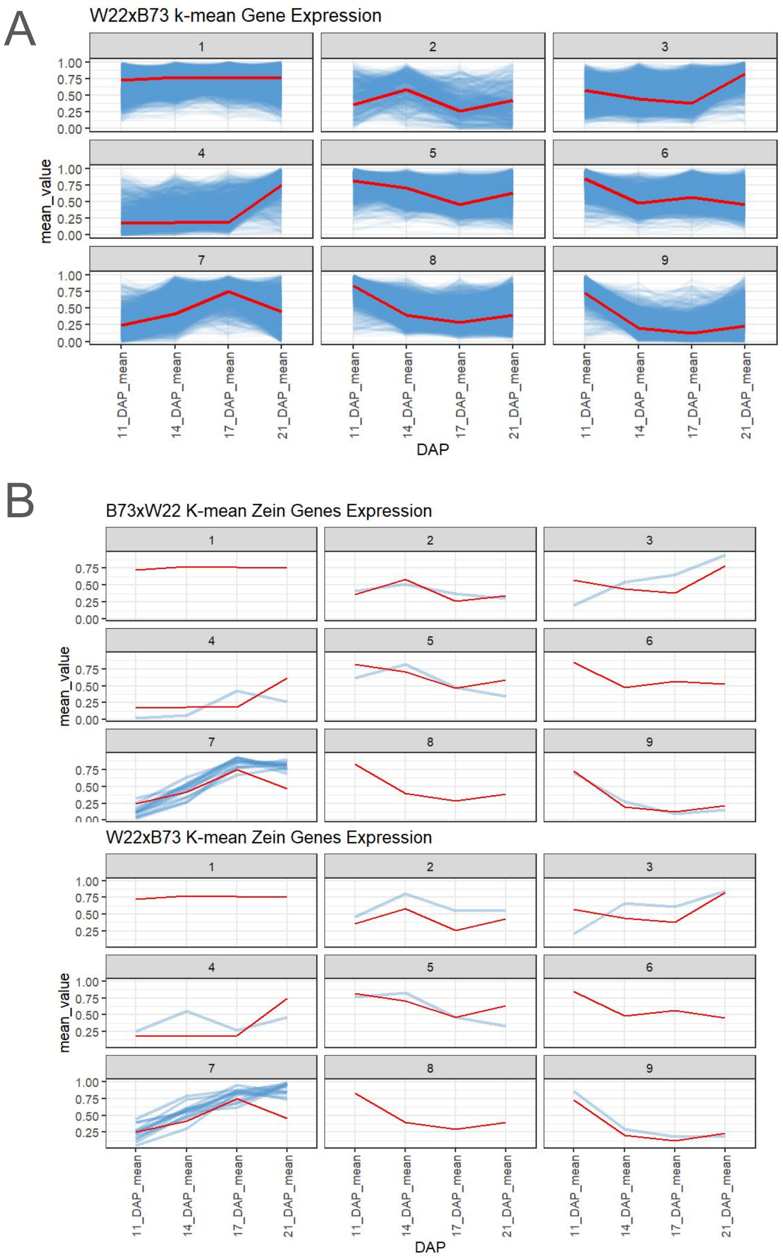


Supplemental Figure 1: kmeans clusters of genes over time. genes were normalized to rpm at each time point divided by max rpm at any time point. Blue lines represent average normalized expression across 3 replicates for each time point. Red lines represent the average of all genes in that cluster in B73. (A) Reads for W22xB73 crosses mapped to B73 genome. (B) Zeins in B73xW22 crosses, and W22xB73 crosses.

### Supplemental figure 2

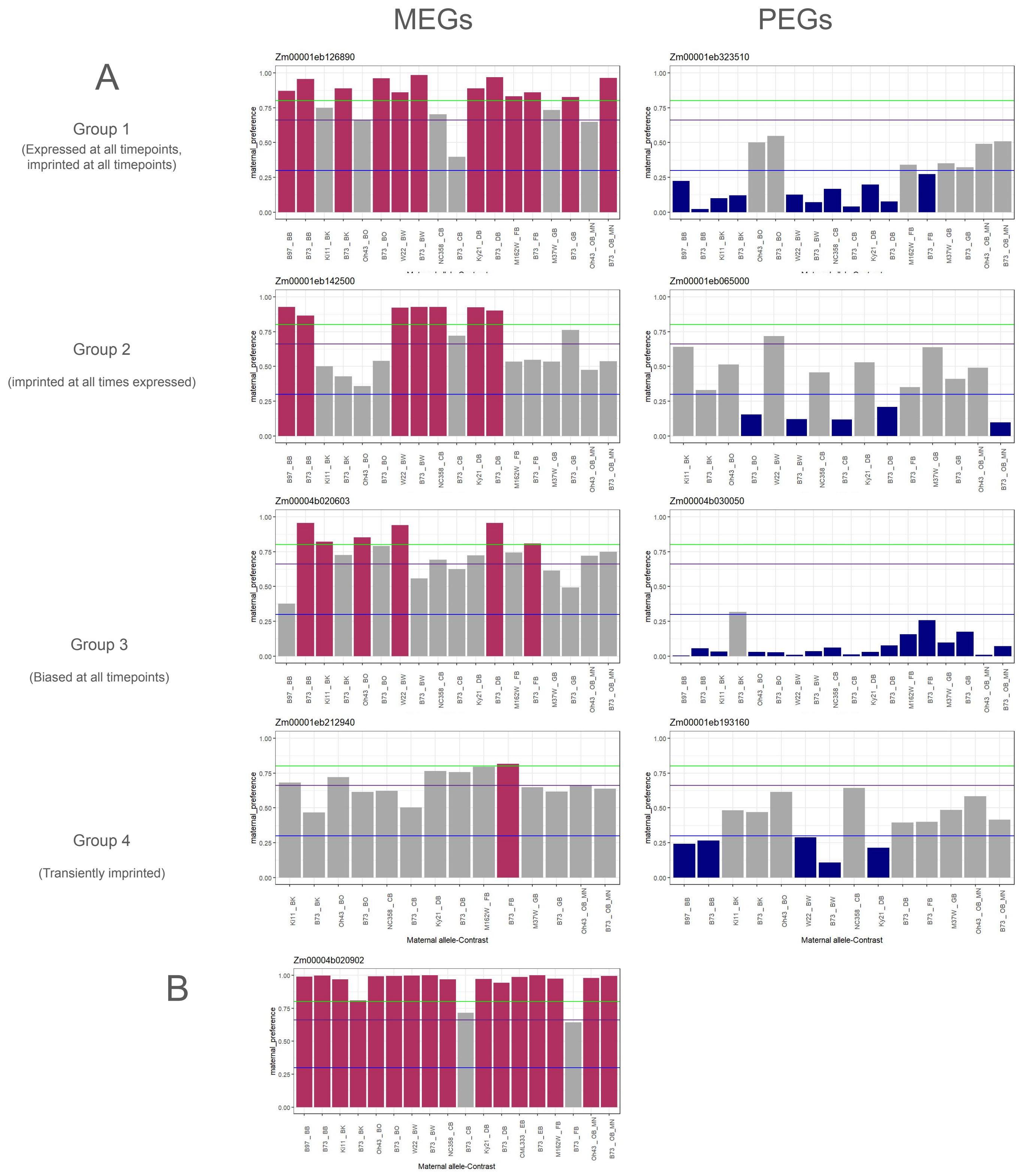


Supplemental Figure 2: Expression patterns compared to maternal preference across lines. Maroon indicates the maternal preference in that cross meets the threshold for RER to be maternally expressed, Blue indicates preference meets the RER threshold to be paternally expressed. Purple horizontal line indicates biparental expression average, green horizontal line indicates maternal RER threshold, blue horizontal line indicates paternal RER threshold. (A) Barplots of maternal preference across lines for example genes of MEGs and PEGs in the four time series groups. (B) Barplot of Fie1 RER across genotypes
